# Farm exposure is associated with human breast milk immune profile and microbiome

**DOI:** 10.1101/2024.10.14.618271

**Authors:** Mary Hannah Swaney, Olivia Rae Steidl, Anastasia Tackett, Samantha Fye, Kristine E. Lee, Irene M. Ong, Casper Bendixsen, Gretchen Spicer, James DeLine, James E Gern, John Lucey, Christine M. Seroogy, Lindsay Kalan, Anne Marie Singh

**Author notes:** Corresponding Authors: Lindsay Kalan, Anne Marie Singh. Contributed equally (Christine M. Seroogy, Lindsay Kalan, Anne Marie Singh).

## Abstract

Prenatal and early life farm exposure, and breastfeeding, are associated with protection from allergic diseases. We hypothesize that farm exposure influences the human breast milk microbiome and immune proteins. The immune protein profiles and microbial communities of 152 human breast milk samples were compared among three maternal farm exposure groups (traditional agrarian, farm, and non-farm) in rural Wisconsin to identify signatures associated with farm status and atopic disease. We found significant differences between farm groups for 23 immune proteins (p-adj<0.05), microbiome diversity (p=2.2E-05), and microbiome richness (p=8.0e-06). Traditional agrarian human breast milk had the highest immune protein levels and microbiome diversity and richness, followed by farm and non-farm human breast milk. Furthermore, Gram-positive bacterial species correlated with IL-23 mediated signaling events (p-adj<1.0E-05). These data suggest that increased farm exposures promotes human breast milk that is more microbially-diverse and rich in immune-associated proteins, ultimately influencing immune development in the infant.

## INTRODUCTION

Atopic diseases, such as atopic dermatitis (AD), asthma, and food allergy (FA) are characterized by a dysregulated immune system that results in allergic inflammation^1–4^. While atopic diseases have strong genetic determinants, their underlying causes cannot be explained by genetics alone. Environmental factors, including increased exposure to sensitizing allergens and reduced stimulation of the immune system by microbial and parasitic factors, are postulated to contribute to atopic disease development, in addition to a complex genetic background^5^. In particular, environmental exposures during early life, which is a critical window in immune development, are likely to strongly influence the risk of atopic diseases. Infant delivery mode, antibiotic use, and tobacco smoke are just a few of many early life exposures associated with an altered risk for the development and severity of allergic conditions^6,7^.

Human breast milk fully satisfies infant nutritional requirements and supports immune development, gut microbiota colonization, and infant growth. Containing a vast number of bioactive molecules that include proteins, oligosaccharides, and minerals, human breast milk is a heterogenous food source whose composition changes in response to the infant’s needs and varies from mother to mother^8–11^. Several observational studies have investigated the role of breastfeeding in protecting the infant from developing atopic diseases. Cytokines, IgA, immune cells, and a unique microbial community in human breast milk, all of which are important factors in immune regulation, may contribute to immunity^11–14^. Indeed, human breast milk IgA levels have been shown to be inversely associated with atopic dermatitis, suggesting a protective effect of human breast milk against atopic disease development^13^. However, findings are inconsistent and some studies have found little to no association between breastfeeding and the development of allergic disease^15–17^. However, these discrepancies may be the result of limited clinical data, the heterogeneous expression of atopic disease, and the high variation seen in human breast milk composition of certain populations of mothers.

Epidemiological and cross-sectional studies have identified an inverse relationship between farm living in early life and the development of atopic diseases including asthma, hay fever, AD, and allergic sensitization^18–21^. Similarly, a traditional agrarian lifestyle, which is practiced by farming-centered, rural societies such as the Plain communities (Amish and Old Order Mennonites), is highly protective against allergic disease^22–25^, and can modulate human breast milk immune and metabolic factors^26^. Furthermore, among farm families, AD prevalence is inversely related to exclusive breastfeeding^27^. Taken together, a possible explanation for these observations may be rooted in differences in farm-related exposures, which may influence human breast milk immune factors and the human breast milk microbiota that could influence infant outcomes.

In this study, we hypothesized that farm exposures are associated with the human breast milk microbiome and immune proteins. To test this hypothesis, we profiled 116 different immune proteins and the microbiota in human breast milk from three rural Wisconsin farm exposure groups: traditional agrarian, farm, and non-farm. We then compared human breast milk immunological and microbial signatures to AD and FA prevalence. Finally, we integrated the immune protein and microbiome datasets to uncover potential associations between the microbiota and immune profile.

## METHODS

### Study Participants

The participants are from an ongoing prospective observational birth cohort study in rural Wisconsin designed to investigate early life farm exposure influence on immune development and respiratory allergies and illnesses in children^27,28^. Participants in the ongoing study were born between 2013-2020. Participants born between 2013-Jan 2018 were enrolled at one study site, Marshfield Clinic. Starting in February 2018, the farm, non-farm, and Traditional Agrarian families were recruited concurrently at two study sites, Marshfield Clinic and LaFarge Medical Clinic (part of Vernon Memorial Healthcare). The study groups are: Farm children (n=63) were defined as residence on a farm or mother working full-time on a farm at the time of pregnant mother enrollment with “Farm” defined as dairy cow or cattle operation. Non-farm mothers/children (n=59) do not reside or work on a farm, or reside within 1/8 of mile for a farm. Traditional Agrarian participants (n=30) are self-declared Amish community members. Human breast milk samples from one TA and 2 non-farm women did not pass sequencing and therefore are not included in the microbiome analyses. Human breast milk samples from n=30 TA women, n=23 women living in farming environments and n=35 living in rural/non-farming environments passed QC for protein analysis were included in protein studies. Written informed consent was obtained from the pregnant mother during the second or third trimester of pregnancy. Study procedures for enrolled participants include questionnaires, environmental assessments, and samples of blood and varied biospecimens for microbiome assessments using centralized collection kits and standardized operating procedures as described in^27,28^. The study was approved by the institutional review boards at the Marshfield Clinic and the University of Wisconsin.

### Farm score calculation

At the prenatal, 2-month and 9-month visits, questions were asked to assess the mother’s and child’s exposure to farm animals and forage. Amount of exposure was scored on a scale of 0 (no contact) to 1 (daily contact) for each animal and summed across the animals (cattle & forage, goats, pigs, poultry, sheep and horses). The farm score of the infant at the 2-month visit was selected for use in analysis.

### Atopic dermatitis and food allergy definitions

Detailed information on questionnaire design has been previously reported^28^. One parent of each child responded to repeat questions about diet, exposures, and early life allergy symptoms (atopic dermatitis, food allergy) during the child’s first two years of life (timepoints: 2 months, 1 year and 2 years). Atopic dermatitis was defined as a positive parent report (“provider told you your child had Eczema (atopic dermatitis, or AD). Food allergy (FA) was defined as positive parent report of a reaction within 2 hours of food ingestion which included skin reactions for typical IgE-mediated foods or involved multiple organ systems and elimination of the food from the child’s diet. Clinical outcomes for this study were determined for participant’s who had consistently answered “yes” throughout the first two years life for AD, and throughout the first 2 years of life for FA. Statistically significant differences across farm status groups for select questionnaire variables (Table 1 and 2) were computed using chisq.test() in R.

**Table 1.**
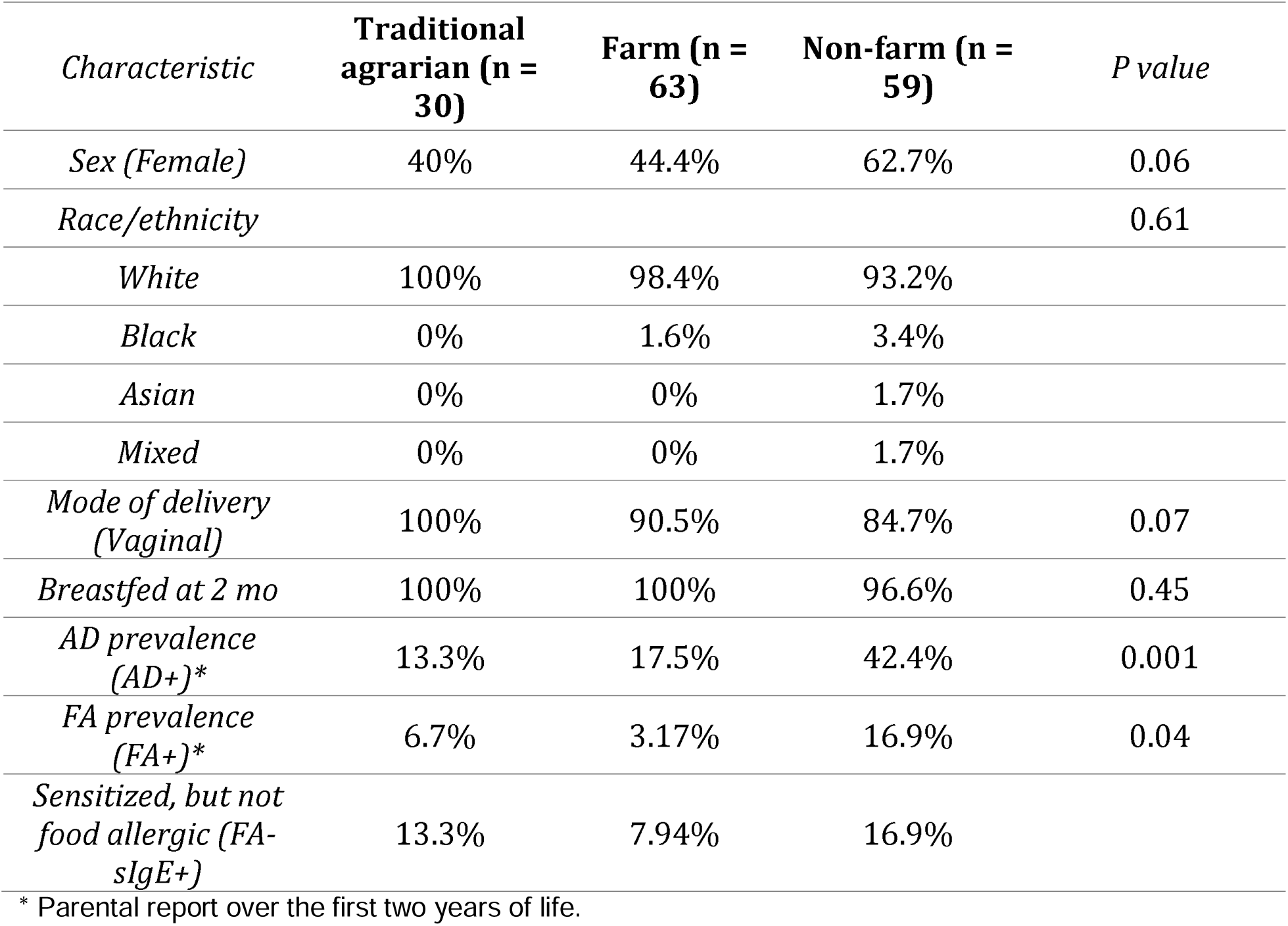
Child Characteristics According to Farm Status.

Infants who were sensitized to a food but without clinical reactivity (ie sensitized but not allergic, or tolerating ingestion of that food despite a positive immunocap test to egg, milk or peanut) are designated as FA-sIgE+. Of note, immunocap testing was only performed to egg, milk and peanut.

### Human breast milk collection and processing

Maternal breast milk was collected from enrolled breastfeeding mothers during scheduled home visits when the child was two months of age (study visit window: 6 weeks to 4 months). Expressed human breast milk followed a standardized protocol adapted from published birth cohort study^44^. The breast was not cleaned before sample collection, therefore collection of some maternal skin microbiota is possible. However, this is likely to be representative of the true microbiota ingested by the infant. Human breast milk was collected into a lot-tracked, sterile container for all study participants. The timing, with respect to time of day or whether foremilk or hindmilk was included in the collection, was not standardized. Samples were transported on ice for up to 4-6 hours, and then transferred to -80°C for long term storage. Human breast milk samples collected within the traditional agrarian community were stored at -20°C for up to two weeks before being transferred to -80°C for long term storage.

Human breast milk samples selected for microbiome analysis represent samples from the study groups (TA n=29, Farm n=63, Non-Farm n=57). Samples were processed as described by Tobin et al.^45^. Briefly, human breast milk was thawed slowly on ice and 5 ml was centrifuged for 10 minutes at 10,000 x g at 4°C to generate a cell pellet. The top lipid/fat layer was removed with a sterile spatula and any residual lipid was removed by pipette. The clear aqueous supernatant was removed and stored in aliquots at -80°C for proteomic analyses. The undisturbed cell pellet was washed with 1 ml of sterile phosphate buffered saline (PBS) and stored at -80°C until DNA isolation. A negative control sample was prepared using PBS and lot-tracked sterile collection supplies; a bacterial community mock control was prepared using the ZymoBIOMICS Microbial Community Standard (Zymo Research, Irvine, California, USA). Both negative and mock controls were processed with study specimens.

### Human breast milk IgA measurement by enzyme-linked immunosorbent assay

IgA levels in human breast milk whey were quantified using a commercial ELISA kit following the manufacturer’s recommendations (Bethyl Laboratories, Montgomery, Texas, USA). Whey samples were assayed in duplicate at a dilution of 1:10,000. Absorbances were read on a BioTek Synergy HTX plate reader at 450 nm. If the %CV of the duplicates was greater than 10%, samples were re-assayed. A standard curve was generated by fitting a 4PL equation to standard dilution series absorbances. Sample concentrations were quantified by interpolating their absorbance from the standard curve generated.

### Sample selection for Olink proteomics

Human breast milk samples for Olink proteomics analysis were selected from the study groups (TA n=30; Farm n=23; Non-farm n=35). For this selection process, a more stringent classification method was used for atopic dermatitis and food allergy status. Control AD-and FA-samples were selected as follows: children with no self-reported atopic dermatitis or food allergy symptoms and with no minor reactions to food over a 2 year period. AD+ samples were selected as follows: repeated self-report of AD over a 6 month period of time (without medication) or self-reported AD with medication prescribed for AD treatment. FA+ samples were selected as follows: self-reported FA and review of healthy history by senior investigators (CMS, AMS) to identify likely cases of food allergy.

### Protein detection and measurement with Olink proteomics

The whey was diluted in a 1:4 ratio to maximize detectable protein levels according to the results of a pilot study. 88 isolated whey human breast milk samples (see above for information on sample selection) were submitted to Olink (Boston, MA), and the inflammation (v.3022) and immuno-oncology (v.3112) 96-plex panels were run on each sample to detect relative levels of 184 total targets. Samples that failed quality control were filtered from analysis (n=1). Additionally, any analyte that had >25% of samples at a non-detectable level was removed, resulting in removal of 54 analytes. Finally, for any analytes that appeared on both panels, only the analyte which had the most detectable data was retained. 116 unique proteins were included after filtering and data cleaning. Any analytes with missing data points (n=1) were estimated by taking the average NPX across all other samples.

### Proteomics statistical analysis

Olink-generated protein expression levels were preprocessed and quality controlled using the Olink NPX Manager software (Olink, Boston, MA) in order to background correct, log2 transform, and normalize samples to a Normalized Protein eXpression (NPX) scale. NPX is an arbitrary relative quantification unit that allows for protein level comparison across samples for a given protein^46^. All figures were generated in R using the OlinkAnalyze (v3.3.0) (REF) and ggplot2 (v3.4.0). Significant differences between two groups were computed using the Welch 2-sample Mann-Whitney U Test at confidence level 0.95 and with Benjamini Hochberg p-value adjustment. Significant differences between three or more groups were computed using the non-parametric Kruskal-Wallis test with a post-hoc Wilcoxon test and Benjamini Hochberg p-value adjustment at confidence level 0.95.

### Human breast milk DNA Isolation

Genomic DNA was isolated using a modified cetyltrimethylammonium bromide (CTAB) buffer protocol^47^. Thawed cell pellets were resuspended with 0.5 ml of 2% CTAB Buffer (Promega Corporation, Madison, Wisconsin, USA), transferred to a Lysing Matrix E tube (MP Biomedicals, Santa Ana, California, USA), vortexed vigorously, and incubated at 65°C for 15 minutes with shaking at 300 rpm. Next, 0.5 ml of phenol:chloroform:isoamyl alcohol (25:24:1) was added and samples were bead beaten using a TissueLyser LT (Qiagen, Hilden, Germany) for 10 minutes at 25 Hz. After centrifugation at 16,000 x g at 4°C, the supernatant was transferred to a heavy phase-lock gel tube (5PRIME, Gaithersburg, Maryland, USA) and the process was repeated starting with the addition of a second volume of CTAB Buffer. An equal volume of chloroform was added to each tube. The tubes were shaken by hand and centrifuged at 12,000 x g at 4°C for 5 minutes. The aqueous layer was transferred to a DNA LoBind Tube (Eppendorf, Hamburg, Germany) containing two volumes of 30% PEG-NaCl precipitation buffer, vortexed vigorously, and incubated for 2 hours at room temperature. Following incubation, samples were centrifuged at 16,000 x g at 4°C for 10 minutes and the supernatant was discarded. The crude gDNA pellet was washed with ice-cold 70% Ethanol and air dried in a biosafety cabinet. gDNA pellets were rehydrated with TE Buffer and stored at -80 C.

### 16S microbiome sequencing and quality control

Purified gDNA was submitted to the University of Wisconsin-Madison Biotechnology Center. DNA concentration was verified fluorometrically. Samples were prepared in a similar process to the one described in Illumina’s 16S Metagenomic Sequencing Library Preparation Protocol, Part # 15044223 Rev. B (Illumina Inc., San Diego, California, USA) with the following modifications: The 16S rRNA gene V4 variable region was amplified with fusion primers (forward primer 515f: 5’-ACACTCTTTCCCTACACGACGCTCTTCCGATCT(N)3/GTGCCAGCMGCCGCGGTAA-3’, reverse primer 806r: 5’-GTGACTGGAGTTCAGACGTGTGCTCTTCCGATCT(N)3/GGACTACHVGGGTWTCTAAT-3’). Region specific primers were previously described in Caporaso et al., 2012 (underlined sequences above)^48^, and were modified to add 3-6 random nucleotides ((N)3/6) and Illumina adapter overhang nucleotide sequences 5’ of the geneLspecific sequences. Following initial amplification, reactions were cleaned using AxyPrep Mag PCR clean-up beads (Axygen Biosciences, Union City, CA). In a subsequent PCR, Illumina dual indexes and sequencing adapters were added. Following PCR, reactions were cleaned using AxyPrep Mag PCR clean-up beads (Axygen Biosciences). Quality and quantity of the finished libraries were assessed using an Agilent DNA 1000 kit (Agilent Technologies, Santa Clara, CA) and Qubit® dsDNA HS Assay Kit (ThermoFisher Scientific), respectively. Libraries were pooled in an equimolar fashion and appropriately diluted prior to sequencing. Paired end 2 x 250 bp sequencing was performed using Illumina MiSeq with the MiSeq Reagent Kit v3 (600-cycle). Images were analyzed using the standard Illumina Pipeline, version 1.8.2.

Quality filtering of the raw reads was performed using the QIIME2 v2023.2.0 quality-control workflow to denoise and filter chimeras of paired-end sequences^49^ along with assembly of reads into amplicon sequencing variants (ASVs) via DADA2^50^. Reads were trimmed and truncated at 12 and 250 nt for forward reads and 20 and 145 nt for reverse reads. We obtained 152 samples and 31.1 million quality-filtered, paired-end reads, with a median of 187,712 paired-end reads per human breast milk sample. Three samples (2190,1780, and 7281) were excluded because of drastically low/high read counts compared to other samples. Amplicon sequencing variants (ASVs) were taxonomically classified using a Naive Bayes classifier based on the sklearn method, which was trained on the Greengenes 2 full-length database^51^. Two mock community standards (Zymo), two negative DNA extraction controls, and one PCR control were also sequenced and processed to ensure a high-quality sequence set. Samples were decontaminated using the DNA extraction and PCR controls with SCRuB v0.0.1^52^.

### Microbiome statistical analysis

All analyses were conducted in the R statistical programming language v4.2.2. To filter rare and contaminant features, ASVs classified as chloroplast or mitochondria and ASVs present at < 0.01% average relative abundance across samples were removed. Total observed ASVs were calculated using the vegan v2.5-7 package and Shannon diversity was calculated using the microbiome v1.12.0 package. For beta diversity analysis, the Bray-Curtis dissimilarity index was calculated for samples using vegdist() from the vegan package. The indices were ordinated using the vegan metaMDS() program for nonmetric multidimensional scaling (NMDS) or the ape v5.4-1 pcoa() program for principal coordinates analysis (PCoA). For differential abundance testing, the Maaslin2 v1.4.0 R package was used^53^, employing the LM analysis method on TSS-normalized and log-transformed relative abundances, with traditional agrarian samples as the reference group.

Human breast milk samples that had both microbiome and protein data available (N=85) were used for partial least squares (PLS) analysis using the mixOmics v6.19.4 R package^54^. The microbiome abundance table was first filtered to include taxa > 0.01% relative abundance and present in 3 or more samples. Abundances were then normalized using cumulative sum scaling (CSS) with the metagenomeSeq v1.32.0 package^55^, followed by log2 transformation. The mixOmics program pls() was run on microbiome and protein data using the canonical framework, which assumes no a priori relationship between the two datasets. Plots were generated using the mixOmics package.

### Pathway Over-representation Analysis

Pathway over-representation analysis (ORA) was performed using the InnateDB Pathway Analysis webform (www.innatedb.com)^56^ using the hypergeometric algorithm and Benjamini Hochberg p-value correction.

## RESULTS

### Study population characteristics

Overall enrollment included mothers from traditional agrarian (TA)communities (n=30), farms (n=63), and rural, non-farm homes (n=59 (Table 1 and Table 2). Human breast milk samples from 1 TA mother and 2 nonfarm mothers failed 16S sequencing. Therefore, we compared the human breast milk of mothers from traditional agrarian communities (n=29), farms (n=63), and rural, non-farm homes (n=57 for scientific analyses. Farm and non-farm participants were recruited as a part of the previously described Wisconsin Infant Study Cohort (WISC) study^27,28^, and the traditional agrarian participants were recruited starting in 2018. The analyses in this study focused on farm status and farm exposure score through child year 1 and AD/FA status through child year 2. Child AD prevalence at age 2 was 13.3% in the traditional agrarian group, 17.5% in the farm group, and 42.4% in the non-farm group. Child FA incidence was lowest in the farm group (3.17%), followed by traditional agrarian (6.7%) and non-farm (16.9%) (Table 1). For maternal incidence of AD, hay fever, and asthma, rates increased for each of these characteristics from the traditional agrarian to farm to non-farm groups (Table 2). A farm score based on frequency of exposure with cattle & forage, goats, pigs, poultry, and horses was calculated for the infant participants at the time of human breast milk collection (2 months). As expected, the farm score of infants from the traditional agrarian group was significantly higher than the non-farm group (p=1.8E-13) (Supplemental Figure 1). Similarly, the score for the farm group was significantly higher than the non-farm group (p=2.1E-12), indicating that the infants living in traditional agrarian and farming environments are coming into more contact with farm animals and plant material.

**Table 2.** Maternal Characteristics According to Farm Status.

| *Characteristic* | **Traditional**<br>**agrarian (n =**<br>**30)** | **Farm (n = 63)** | **Non-farm (n**<br>**= 59)** | *P-value* |
| --- | --- | --- | --- | --- |
| *Self-reported AD* | 13.3% | 20.6% | 23.7% | 0.51 |
| *Self-reported hay*<br>*fever* | 30% | 31.7% | 37.3% | 0.73 |
| *Self-reported*<br>*asthma* | 0% | 20.6% | 32.2% | 0.002 |
| *Consumption of*<br>*raw farm milk*<br>*during*<br>*pregnancy* | 96.7% | 7.9% | 0% | <2.2E-16 |

### Human Breast milk protein profiling

To assess proteomic differences in human breast milk collected from mothers with varying farm exposure, cytokines, chemokines, and other immune-related proteins were measured using two Olink panels focused on immune responses and inflammation (Olink, Boston, MA). When comparing among traditional agrarian, farm, and non-farm mothers, there were significant differences for 23 proteins between the groups, determined using a multi-group Kruskal-Wallis test (traditional agrarian vs farm vs non-farm) (Figure 1A). These human breast milk proteins differed most between traditional agrarian and non-farm mothers, and over half were significantly different between traditional agrarian and farm samples. Two proteins (TWEAK and TNFRSF4) differed between farm and non-farm human breast milk samples and were not significantly different between the traditional agrarian and farm. Overall, human breast milk from mothers living in traditional agrarian communities contained the highest median levels of these proteins, while levels were more similar across the farm and non-farm breast milk. Individual protein levels were associated with farm score for many of the proteins (Figure 1B). Furthermore, when hierarchical clustering of the 23 significantly-different proteins was completed, one cluster principally comprised of traditional agrarian samples, while farm samples tend to form inter-mixed clusters with non-farm and traditional agrarian samples (Figure 1C). The variance within the farm group could be a result of differences in farm-related exposures among farm mothers (Supplemental Figure 1).

**Figure 1.**
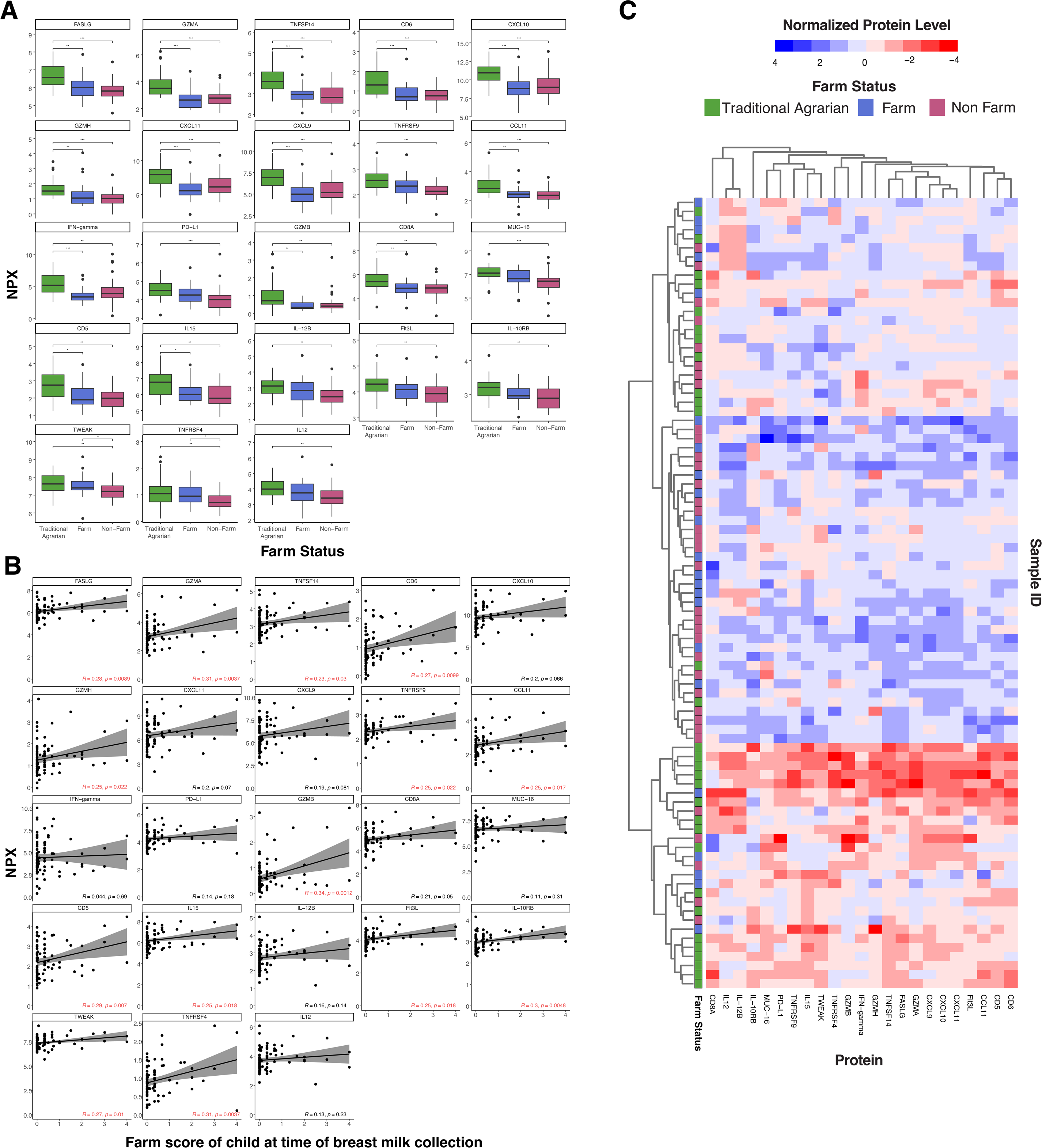
Human breast milk from traditional agrarian mothers exhibits high immunologic protein levels. The relative levels of 116 immune proteins were measured using the Olink platform and compared between the human breast milk from traditional agrarian, farm, and non-farm mothers (TA n=30; Farm n=23; Non-farm n=35), with the 23 statistically significant proteins shown here. Relative levels are shown in the Normalized Protein eXpression (NPX) scale, which is an arbitrary relative quantification unit. Statistically significant differences between groups were calculated using a Kruskal-Wallis test, followed by a Dunn posthoc test with Benjamini-Hochberg p-value adjustment. * p<0.05, ** p<0.01, *** p<0.001. B) Pearson correlation was performed using the 23 statistically significant proteins and NPX values. Black lines represent the linear regression line and the shaded gray area represents the 95% confidence interval. Significant correlations are highlighted in red text. C) For all human breast milk samples, the 23 statistically significant proteins and their normalized expression levels (scaled to values between -4 and 4) are displayed as a heatmap for all breast milk samples. Hierarchical clustering was performed on samples and proteins using the complete linkage method. Samples are colored according to farm status.

Over-representation analysis was performed to determine if the 23 significant proteins were related to specific biological pathways. This method assesses statistically whether the set of proteins shares more or fewer proteins with protein pathways than would be expected by chance. Findings of the analysis reveal that many pathways were enriched for these 23 proteins, including cytokine-cytokine receptor interactions, IL12-mediated signaling events, and downstream signaling in naïve CD8+ T cells (p < 1.0E-05) (Supplemental Data Sheet 1). Secretory IgA produced within the mother’s breast milk plays a critical role in immune development, protection against infection and inflammation, and development of the gut microbiome^29^. We thus compared the concentration of breastmilk IgA among the three farm groups, and found significantly higher levels in traditional agrarian human breast milk compared to farm (p<0.05) and non-farm (p<0.05) (Supplemental Figure 2). There was not a significant difference in IgA between farm and non-farm samples.

### Human Breast milk immune protein levels and atopic disease status

We next evaluated if differences in human breast milk immune proteins were associated with the expression of atopic disease (consistent expression of atopic dermatitis and food allergy during the first two years of life). Human breast milk levels of TIE2 and CD27 were nominally lower in mothers who had infants with AD, but these differences were non-significant after adjusting for multiple comparisons (Figure 2A). Similarly, levels of 9 proteins (4E-BP1, ADA, ARG1, AXIN1, CD83, Gal-1, GZMA, IL18, and ST1A1) were significantly different between mothers who had infants with or without FA or were FA-sIgE+, (multi-group Kruskal-Wallis FA+ vs FA-sIgE+, FA-) but were not significant after adjusting for multiple comparisons (p-value > 0.05) (Figure 2B). Interestingly, of these proteins, the highest levels tended to be found within samples.

**Figure 2.**
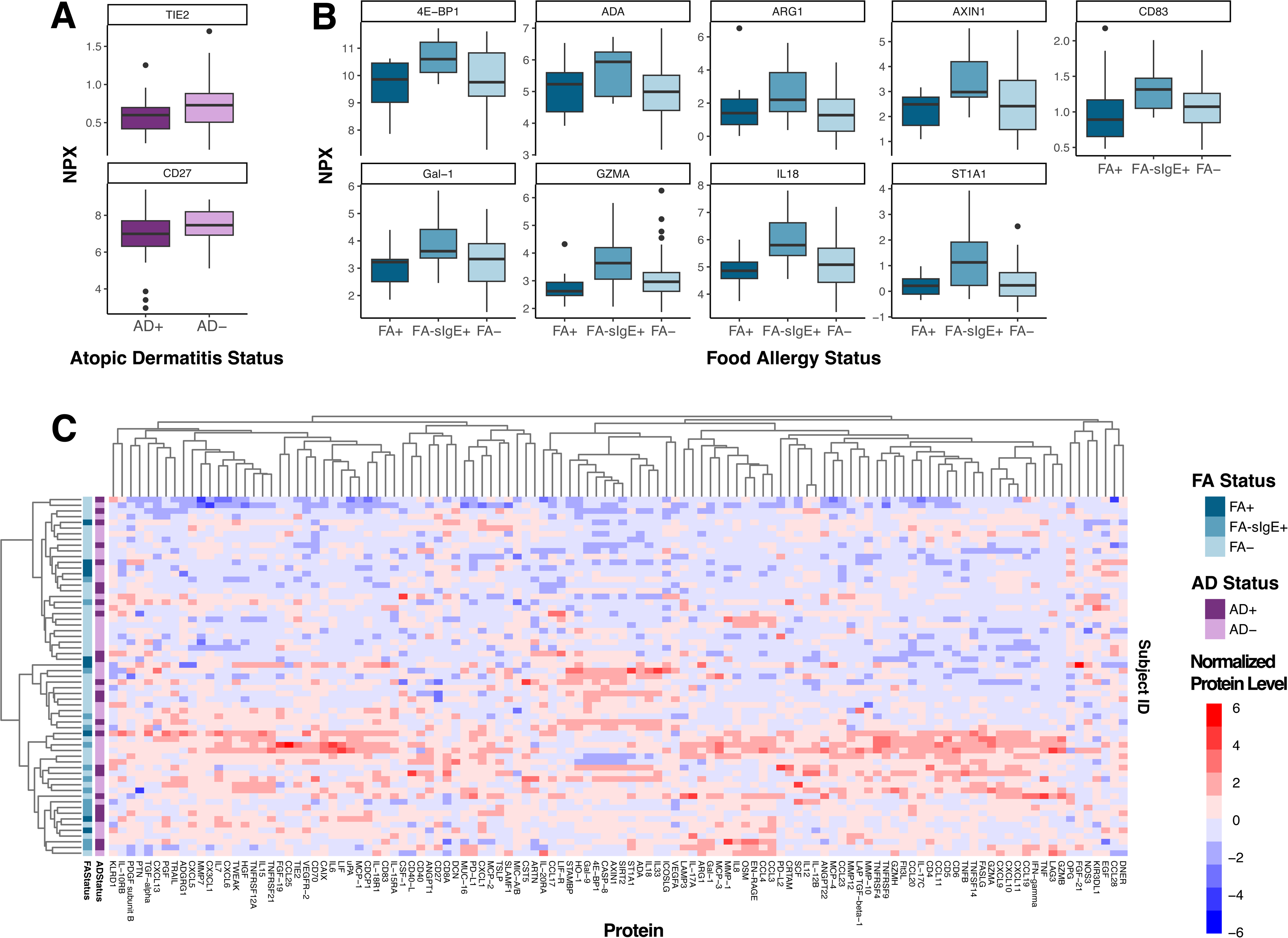
Human breast milk immunologic protein levels and infant atopic disease status. **A)** Proteins that were statistically significant before p-value correction across human breast milk samples from infants with atopic dermatitis (AD) (n=40) and without AD (n=79). **B)** Proteins that were statistically significant before p-value correction across human breast milk samples from infants with food allergy (FA) (n=14), infants who sensitized but not allergic (FA-sIgE+ (n=19), and infants without FA (n=81), determined using a Kruskal-Wallis test (FA+ vs FA-sIgE+. vs FA-). **C)** For all samples with an assignment for both AD and FA status (n=63), all 116 immunologic proteins and their normalized expression levels (scaled to values between -6 and 6) are displayed as a heatmap. Hierarchical clustering was performed on samples and proteins using the complete linkage method. Samples are colored according to AD status and FA status.

### Composition of the human breast milk microbiome

We determined whether the human breast milk microbiome composition differed between the traditional agrarian, farm, and non-farm exposure groups. We performed 16S rRNA sequencing of 117 human breast milk samples. After quality filtering, we observed 2,639 unique amplicon sequence variants (ASVs), a proxy for bacterial species, across 37 bacterial phyla in all samples. Among these phyla, Pseudomonadota (formerly Proteobacteria, 46.9%) accounted for the largest average proportion of the community, followed by Bacillota group D (formerly Firmicutes, 43.4%), Actinomycetota (formerly Actinobacteria, 4.3%), and Bacteroidota (formerly Bacteroidetes, 2.2%) (Supplemental Figure 3). Other phyla represented an average 0.33% of the human breast milk microbial community. At the genus level, human breast milk was largely dominated by *Streptococcus* (24.5%), *Acinetobacter* (23.1%), *Staphylococcus* (14.4%), and *Pseudomonas* (5.8%). Present at an average relative abundance below 5% included *Veillonella, Corynebacterium, Gemella,* and *Rothia*. Other genera were present at an average relative abundance below 1%, accounting for around 19.3% of the microbial community composition on average (Supplemental Figure 3).

Alpha diversity of each sample, defined as the within-sample microbial diversity, was calculated with respect to ASV richness and evenness using the Shannon diversity index and total observed ASVs. When considering sex of the infant, both Shannon diversity and total observed ASVs were higher in the breast milk of mothers with male infants as compared to female infants (p=0.0089 and p=0.0065, respectively) (Figure 3A,B). When assessing beta diversity, or the similarity of microbial communities between samples, composition of bacteria communities were not significantly different between mothers with male versus female infants (PERMANOVA p-value = ns) (Figure 3C). Human breast milk nutrient composition has previously been shown to be different between mothers with male versus female babies, with that from mothers of male infants being higher in fat and energy content ^30,31^, providing a possible explanation for the observed differences in human breast milk microbiota diversity.

**Figure 3.**
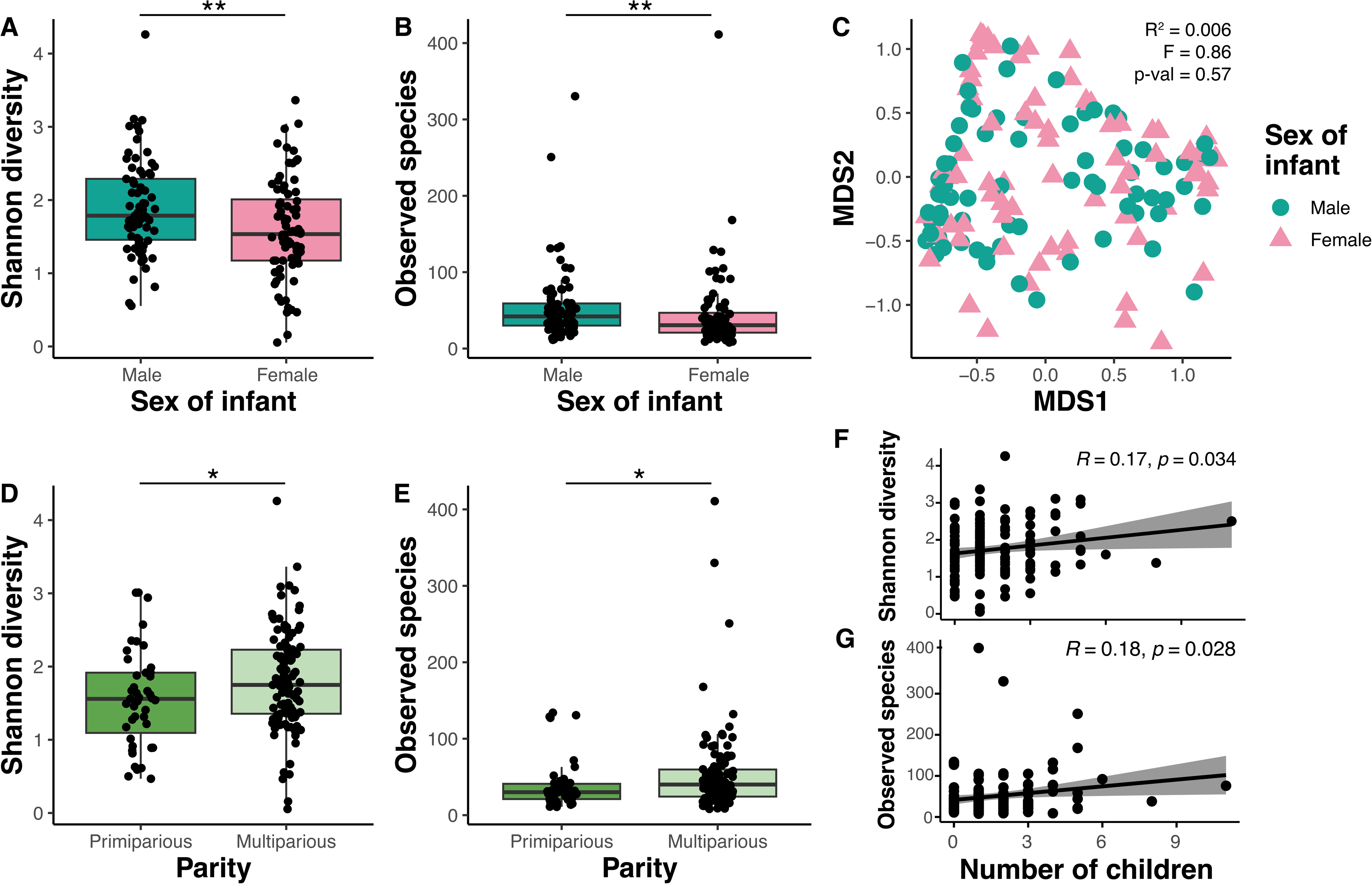
Human breast milk microbiome diversity is associated with sex of the infant and maternal parity. **A)** The Shannon diversity index and **B)** total observed amplicon sequencing variants (ASVs) were calculated for each sample and grouped by sex of the infant (Male n=73, Female n=76). **C)** Non-metric dimensional scaling (NMDS) plot based on the distance matrix of Bray-Curtis dissimilarity of the microbial communities across samples. PERMANOVA was performed to assess statistical significance of microbial community composition differences between infant sexes. **D)** Shannon diversity index and **E)** ASVs of each sample, grouped by maternal parity (Primiparous n=43, Multiparous m=106). **F)** Pearson correlation was performed using **F)** Shannon diversity index or **G)** ASVs and the number of other children the mother has given birth to. Black lines represent the linear regression line and the shaded gray area represents the 95% confidence interval.

Both Shannon diversity and observed ASVs were significantly higher in multiparous mothers (p=0.043 and p=0.013, respectively) (Figure 3D,E). Also, there was a significant positive correlation between the number of children the mother had previously given birth to and both Shannon diversity (p=0.034) and observed ASVs (p=0.028) (Figure 3F,G).

### Human breast milk microbiome differs by farm status

Shannon diversity was significantly higher in the breast milk of the traditional agrarian group compared to both farm (p=0.010) and non-farm (p=1.1E-05) groups (Figure 4A). Similarly, total observed ASVs was significantly higher in the traditional agrarian group compared to both farm (p=0.001) and non-farm (p=4.2E-06) groups (Figure 4B).

**Figure 4.**
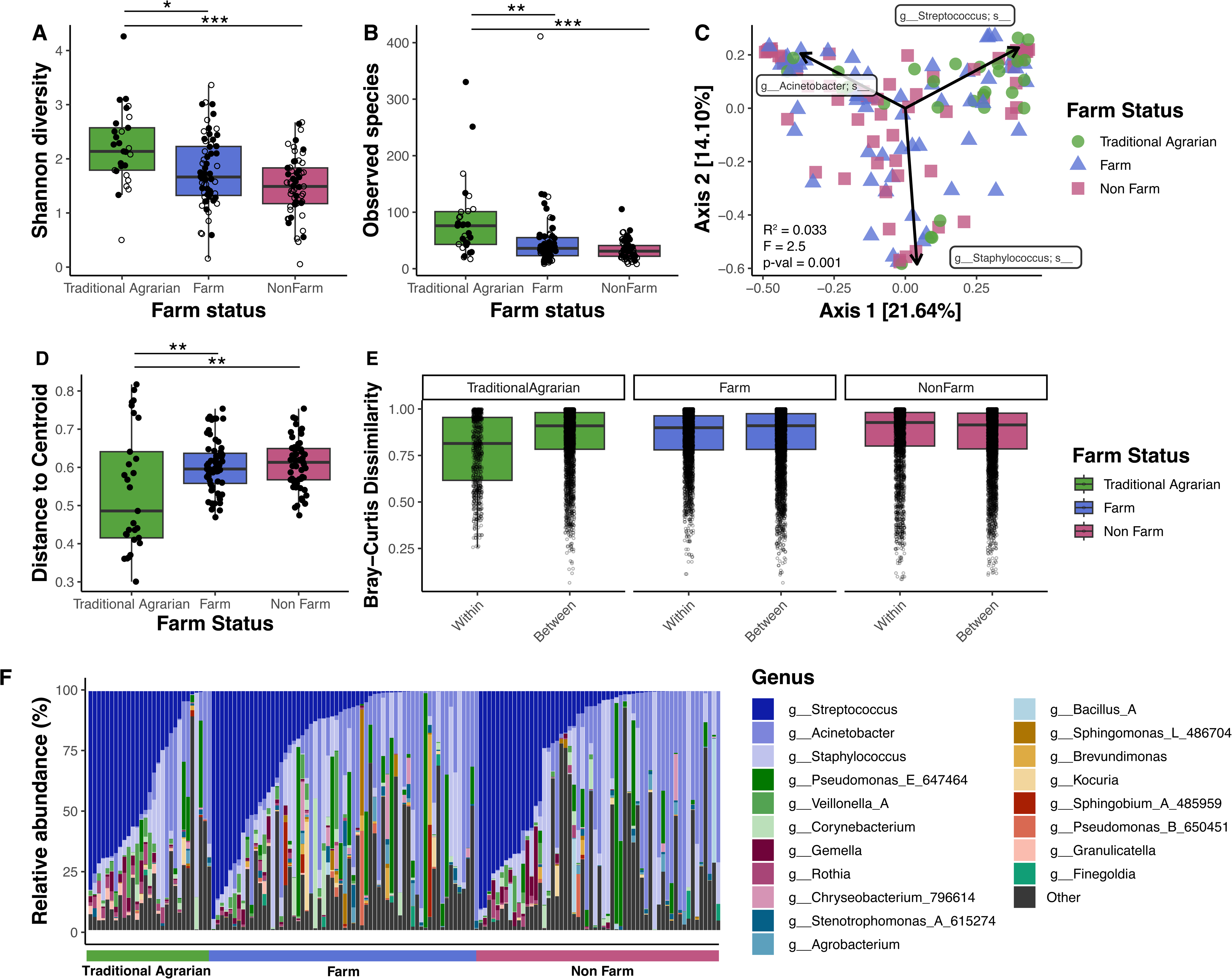
Breast milk microbiome diversity and composition is significantly different among mothers with varying farm-related exposure. 16S rRNA sequencing was performed on DNA extracted from human breast milk samples (TA n=29, Farm n=63, Non-Farm n=57). **A)** The Shannon diversity index and **B)** total observed amplicon sequence variants (ASVs) were calculated for each sample, grouped by farm status. **C)** Principal coordinate analysis (PCoA) biplot based on the distance matrix of Bray-Curtis dissimilarity of the microbial communities across samples. The top three ASV loadings are represented as arrows. PERMANOVA was performed to assess statistical significance of microbial community composition differences between farm status. **D)** Beta-dispersion values (distance from the centroid) for each sample, grouped by farm status. **E)** Bray-Curtis dissimilarity values for all samples against all samples within or between farm status group. **F)** Microbial community composition of each sample based on relative abundance of ASVs at the genus level. Samples are ordered by farm status and Streptococcus abundance. “Other” reflects genera apart from those shown in the legend and also ASVs that did not have a genus assignment. p<0.05, **p<0.01, ***p<0.001 based on a Kruskal-Wallis test, followed by a Dunn posthoc test with Benjamini-Hochberg p-value adjustment.

Farm exposure was also related to microbiome composition (PERMANOVA p-value = 0.001) (Figure 4C). Traditional agrarian samples showed significantly different dispersion from both the farm and non-farm samples (p=0.008 and p=0.003, respectively) (Figure 4D) and exhibited the lowest median within-group Bray-Curtis dissimilarity when compared to that of the farm and non-farm groups (Figure 4E). This suggests that there is less variation and more similar composition of the microbial communities found in traditional agrarian breast milk when compared to farm or non-farm samples. Indeed, high inter-sample variability is observed in the microbial community composition of breast milk from farm and non-farm mothers, while traditional agrarian samples had more similar community composition at the relative abundance level, generally dominated by Bacillota including *Streptococcus* and *Staphylococcus* (Figure 4E, Supplemental Figure 4A,B). Conversely, farm and non-farm microbiomes are more frequently dominated by Pseudomonadota and exhibit more variable community compositions across samples. At the ASV level, 10 ASVs are present at significantly higher relative abundance in breast milk from traditional agrarian mothers as compared to farm and non-farm mothers, with the most significantly different being ASVs from the *Rothia, Granulicatella*, *Streptococcus*, *Paulgensenia*, and Veiillonellaceae taxa q-value < 0.05) (Supplemental Figure 4C).

### Relationship between immunologic proteins and the human breast milk microbiome

Human breast milk is a complex biological fluid where resident microbes likely stimulate host immune responses and immune activity modulates microbial populations. To provide insight into pathways and/or interactions between the breast milk microbiota and the local immune environment, we employed the partial least squares (PLS) method to interrogate these relationships using 85 human breast milk samples with both microbiome and immunological protein data. The correlation structure between ASVs and immune proteins is displayed in Figure 5A, revealing distinct clusters of ASVs that correlate with sets of proteins. For example, cluster A4 reveals strong positive correlations between numerous proteins and Gram positive ASVs from the Actinomycetota and Bacillota phyla. Conversely, cluster D1 exhibits generally negative correlations between Gram negative ASVs from the Pseudomonadota phylum and certain proteins. The strongest correlations from the heatmap (values greater than 0.5) are presented as a relevance network in Figure 5B, which reveals a cluster composed of all positively-correlated proteins and ASVs4 (Figure 5B). Notably, this cluster consists of proteins that contribute to antimicrobial and antiviral innate immune responses (e.g., CXCL10, CXCL9, IL-6, IL-17 and others), and over-representation analysis reveals that the top pathways this set of proteins is enriched in are chemokine receptors bind chemokines (REACTOME, p < 1.0E-05), IL-23 mediated signaling events (PID NCI p < 1.0E-05) (e.g., CCL2, CD4, CXCL9, IL18R1, IL6), CXCR3-mediated signaling events (PID NCI p = 2.0E-05) (CCL11, CXCL9, CXCL10, CXCL11), and cytokine-cytokine receptor interactions (KEGG, p < 1.0E-05) (Supplemental Data Sheet 1).

**Figure 5.**
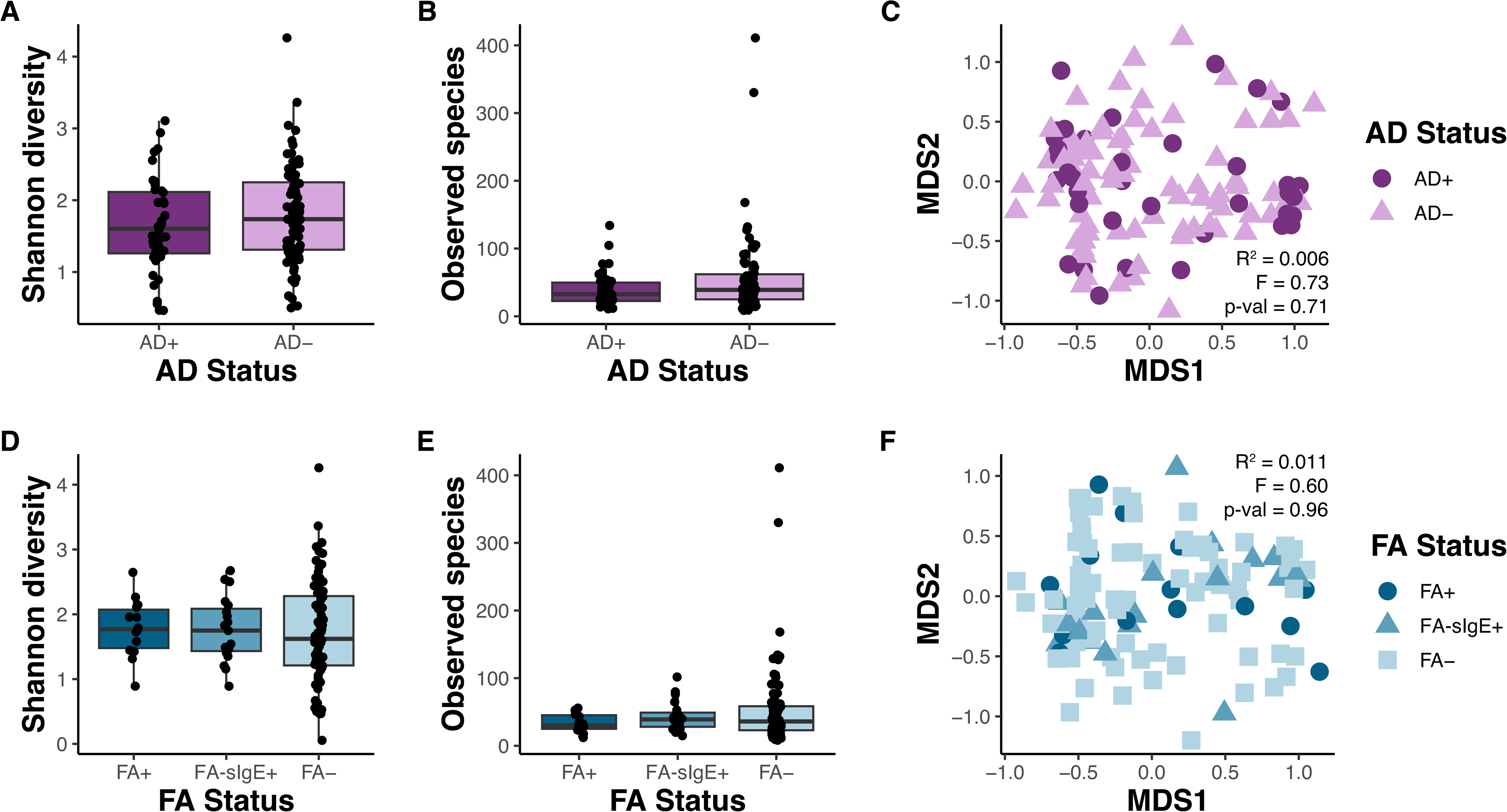
Integration of human breast milk immune protein expression with microbiome composition. PLS canonical correlation analysis was performed using the mixOmics platform to integrate relative protein levels with microbiome abundance data for human breast milk samples (n=85). A) Clustered image map (CIM) representing the correlation structure of the protein and microbiome datasets from the first two latent components. ASVs, colored by taxonomic assignment at the phylum level, represent rows and proteins represent columns. Hierarchical clustering was performed on ASVs and proteins using the complete linkage method. B) Relevance network representing the correlations from A) that are greater than or equal to 0.5. Nodes indicate either protein (green rectangle) or ASV (blue oval colored by phylum) Edges represent correlations between an ASV and protein; red = positive correlation, blue = negative correlation.

### Human breast milk microbiome and atopic disease status

To determine if there were differences in the breast milk microbiome among mothers with infants with or without AD or FA, we compared alpha and beta diversity for AD and FA statuses. No significant differences in the microbial community diversity was observed between AD+ or AD-infants (Figure 6A-B), and microbial community composition did not group by AD status (Figure 6C). However, Shannon diversity tended to be lower in breast milk from mothers with AD+ infants. Furthermore, no significant differences were observed for alpha or beta diversity when considering FA status (Figure 6D-F). While strong differences were not observed in the human breast milk microbiome when considering AD or FA status, underlying variables, such as the degree of farm-related exposure and the heterogeneity of atopic disease, may play a role.

**Figure 6.**
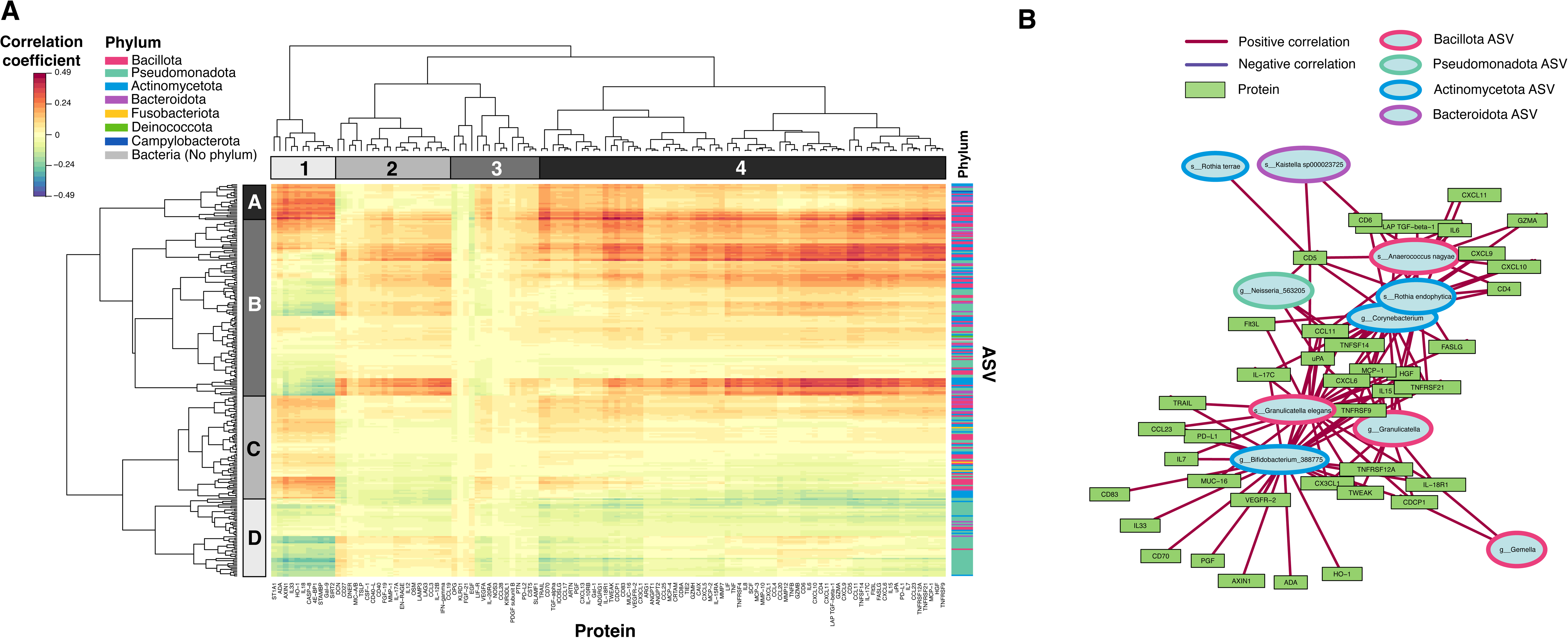
Human breast milk microbiome diversity is not significantly different between mothers of infants with or without atopic disease. A and D) The Shannon diversity index and B and E) total observed amplicon sequence variants (ASVs) were calculated for each sample (AD+ n=40, AD-n=79). C and F) Principal coordinate analysis (PCoA) biplot based on the distance matrix of Bray-Curtis dissimilarity of the microbial communities across samples. Samples are grouped by atopic dermatitis (AD) status in A-C or food allergy (FA) status in D-F (FA+ n=14, RAST+ n=19, FA-n=81).

## DISCUSSION

Prenatal and early life farm exposures, as well as breastfeeding, have been identified to be associated with the farm protective effect against atopic disease development^18–21^. Herein, we compared the immune protein profiles and microbial communities of breast milk from three farm exposure groups in rural Wisconsin to identify potential signatures associated with farm status and early life allergic disease.

Our study revealed that almost all immune proteins were expressed at the highest levels in breast milk from traditional agrarian mothers, as compared to farm and non-farm mothers. This suggests that a traditional agrarian lifestyle may promote a distinct postnatal immune development and gut microbial community. Furthermore, our results show that of the 23 proteins that are significantly different by farm status, 14 of these proteins are positively correlated with farm score, which is a measurement of exposures to animals and other farm-related factors. Peroni et al. also found that farm exposure was associated with higher levels of certain cytokines in human breast milk, notably TGF-β1 and IL-10^32^. Overall, our findings support some farm exposures influence human breast milk cytokine concentrations which may in turn influence infant immune development.

Human breast milk, in addition to containing immunomodulatory cytokines and proteins, also contains a distinct microbial community that is transferred to the infant through breastfeeding and is thought to stimulate and promote maturation of the infant immune system^34,35^. Our findings revealed that breast milk microbiome from traditional agrarian mothers is significantly more diverse and contained more species overall compared to both farm and non-farm breast milk. These findings are in agreement with previous studies of the microbial environment surrounding a traditional agrarian lifestyle, which identified farm exposure to be positively associated with microbial diversity in dust samples^24,36^. This also further supports the hypothesis that influential farm-related exposures may be microbial in nature^36,37^. Our findings suggest that increased farm exposures may promote a more diverse human breast milk microbiome, which is ultimately transferred to the infant upon breastfeeding.

Traditional agrarian breast milk, while having a more diverse microbiome, did not appear to have a microbial community totally unique to that of farm or non-farm breast milk. However, traditional agrarian breast milk samples were more likely to be dominated by taxa from the Actinomycetota and Bacillota phylum and were found to be differentially enriched in select ASVs. This suggests that traditional agrarian breast milk may have a more diverse microbial signature and enriched in certain taxa. This microbial signature could contribute to the beneficial and protective nature conferred by the traditional agrarian lifestyle.

Bacteria in the GI tract can stimulate early immune system development^38^, therefore we investigated the potential relationship between the microbiome and immunological protein profile of the human breast milk within our study. Using data integration of the two datasets, we found that certain bacteria predominantly from the Actinomycetota and Bacillota phyla, which are generally Gram-positive, were strongly correlated with numerous proteins involved in antimicrobial and antiviral innate immune responses. Similarly, select bacteria from the Gram-negative Pseudomonadota phyla formed negative correlations with several proteins, although the functional relationship between these proteins was unclear. This suggests that Gram-negative and Gram-positive bacteria within the human breast milk microbiome may differentially influence the immune landscape of human breast milk, or vice versa. Indeed, fundamental differences in the innate immune responses to Gram-positive and -negative bacteria have been noted previously^39,40^. Interestingly, we also observed that the Gram-positive Actinomycetota and Bacillota phyla tended to be enriched within the breast milk of the traditional agrarian group, while the Gram-negative Pseudomonadota phyla was generally reduced. Taking together the finding that these phyla are differentially associated with different immune proteins, increased farm-related exposures could favor a human breast milk microbiome dominated by the Actinomycetota and Bacillota phyla, which in turn leads to a more protective breast milk immune profile to be transferred to the infant.

Because increased farm exposures are associated with decreased incidence of atopic disease, we investigated if there were differences in human breast milk immune protein levels and the microbiota associated with AD and FA status within our study. We did not detect significant differences across AD or FA status, which may be a result of the heterogeneous nature of atopic disease as well as the current study being underpowered. This is an area of interest that requires future work, and we believe that carrying out a larger, higher-powered study to investigate atopic disease and human breast milk composition would further delineate these patterns.

In summary, this study provides evidence that farm exposure is associated with differences in human breast milk immunological protein levels and the human breast milk microbiome. Because human breast milk is a rich source of bioactive and immunomodulatory molecules as well as microorganisms for infant gut colonization, elucidating the influence of the environmental exposures on human breast milk composition is essential for understanding infant development, health, and risk for atopic disease. Future directions of our study include investigating other factors of human breast milk, such as the metabolome, proteome, and human milk oligosaccharide (HMO) composition, and their relationship with farm exposure, atopic disease, and infant development. Recent work has identified an association between residential green environments with HMO composition^41^, demonstrating that external factors are likely important mediators of human breast milk composition. Furthermore, other studies have begun to shed light on the relationship among the different human breast milk components, including the metabolome and microbiome^42^, as well as human breast milk fatty acid composition and the microbiome^43^. Therefore, integration of multiple biological datasets is needed to further our understanding of human breast milk compositional differences and their ultimate influence on the infant.

## ACKNOWLEDGEMENTS

We thank the study families for their participation in this study. The authors thank the University of Wisconsin Biotechnology Center DNA Sequencing Facility for providing sequencing facilities and services.

## FUNDING

This work was supported by grants from the USDA-National Institute for Food and Agriculture WIS03063 [JL and AMS], National Institutes of Health NIH UG3 OD035509 [AMS], NIAID U19 AI104317 [CMS], CTSA 1UL1TR002373 [CMS], NIAMS F31AR079846 [MHS], Wisconsin Partnership Program [CMS], Canada Research Chair in Skin Microbiome & Infection [LRK]. The content is solely the responsibility of the authors and does not necessarily represent the official views of the National Institutes of Health.

## Sources of Funding

USDA-National Institute for Food and Agriculture WIS03063 [JL and AMS]

NIH/NIAID U19 AI104317 [CMS]

NIH/CTSA 1UL1TR002373 [CMS]

Wisconsin Partnership Program [CMS]

NIAMS F31AR079846 [MHS]

Canada Research Chair in Skin Microbiome & Infection [LRK]

NIH UG3 OD035509 [AMS]

## Author contributions

AMS, CMS, CB, LK, JG, and JD contributed to study conception and design. GS collected samples from participants. SF and ORS prepared samples for protein expression analysis and microbiome sequencing. MHS, ORS, CMS, KL, and IMO performed analyses. AT and JL contributed to overall conceptualization and provided critical input to study design and interpretation. KL provided guidance on statistical considerations. MHS, LK, AMS, CMS, and SF wrote the manuscript with input from all authors.

**Supplemental Figure 1.** Child farm score at the time of human breast milk collection (2 months), which is based on frequency of exposure with cattle & forage, goats, pigs, poultry, sheep, and horses (see Methods), is shown for traditional agrarian, farm, and non-farm mother-infant pairings (TA n=30, Farm n=63, Non-Farm n=59).

**Supplemental Figure 2. Breastmilk samples from TA mothers contain significantly increased IgA compared to Farm and Non-Farm groups.** Breastmilk IgA levels at 2 months are shown compared to farming status.*p<0.05

**Supplemental Figure 3. A)** Average relative abundances of the top phyla present across all 149 human breast milk samples. Phlya present at an average of less than 1% relative abundance are grouped into “Other”. The Bacillota phylum is subdivided into groups A-D because it is polyphyletic within the Greengenes2 reference tree, which was used for Qiime2 taxonomic assignment. **B)** Average relative abundances of the top genera present across all human breast milk samples. Genera present at an average of less than 1% relative abundance are grouped into “Other”.

**Supplemental Figure 4. A)** Microbial community composition of each sample based on relative abundance of ASVs at the phylum level. Samples are grouped by farm status and ordered by the Bacillota (D) phylum. **B)** Violin plot of Bacteroidota, Actinomycetota, Pseudomonadota, and Bacillota relative abundances in each human breast milk sample. *p<0.05, **p<0.01, ***p<0.001 based on a Kruskal-Wallis test, followed by a Dunn posthoc test with Benjamini-Hochberg p-value adjustment. **C)** Differentially abundant ASVs between farm/non-farm and traditional agrarian samples, with traditional agrarian as the reference group, determined using MaAsLin2. ASVs with a corrected q-value less than 0.05 are shown. Color within the heatmap takes into account q-value and sign of the coefficient (effect estimate), where a darker color indicates a larger difference between the test group (Farm or Non-Farm) and the reference group (Traditional Agrarian).

**Supplemental Data S1**. Over Representation Analysis for significantly different cytokine levels by Farm Status (Traditional Agrarian, Farm, Non-Farm)

**Supplemental Data S2**. Over Representation Analysis for cytokines correlated with the breast milk microbiota in the PLS analysis

## Notes

### Competing Interest Statement

The authors have declared no competing interest.

## REFERENCES

1. Gold MS, Kemp AS. Atopic disease in childhood. Med J Aust. 2005;182:298–304.

2. Pellerin L, Jenks JA, Bégin P, Bacchetta R, Nadeau KC. Regulatory T cells and their roles in immune dysregulation and allergy. Immunol Res. 2014;58:358–68.

3. Wang Y, Liu L. Immunological factors, important players in the development of asthma. BMC Immunol. 2024;25:50.

4. Kenney HM, Battaglia J, Herman K, Beck LA. Atopic dermatitis and IgE-mediated food allergy: Common biologic targets for therapy and prevention. Ann Allergy Asthma Immunol. 2024;133:262–77.

5. Novak N, Bieber T. Allergic and nonallergic forms of atopic diseases. J Allergy Clin Immunol. 2003;112:252–62.

6. Gabryszewski SJ, Dudley J, Grundmeier RW, Hill DA. Early-life environmental exposures associate with individual and cumulative allergic morbidity. Pediatr Allergy Immunol. 2021;32:1089–93.

7. Murrison LB, Brandt EB, Myers JB, Hershey GKK. Environmental exposures and mechanisms in allergy and asthma development. J Clin Invest. 2019;129:1504–15.

8. Ballard O, Morrow AL. Human milk composition: nutrients and bioactive factors. Pediatr Clin North Am. 2013;60:49–74.

9. Vass RA, Kemeny A, Dergez T, Ertl T, Reglodi D, Jungling A, et al. Distribution of bioactive factors in human milk samples. Int Breastfeed J. 2019;14:9.

10. Rajani PS, Seppo AE, Järvinen KM. Immunologically Active Components in Human Milk and Development of Atopic Disease, With Emphasis on Food Allergy, in the Pediatric Population. Front Pediatr. 2018;6:218.

11. Notarbartolo V, Giuffrè M, Montante C, Corsello G, Carta M. Composition of Human Breast Milk Microbiota and Its Role in Children’s Health. Pediatr Gastroenterol Hepatol Nutr. 2022;25:194–210.

12. Böttcher MF, Jenmalm MC, Garofalo RP, Björkstén B. Cytokines in breast milk from allergic and nonallergic mothers. Pediatr Res. 2000;47:157–62.

13. Orivuori L, Loss G, Roduit C, Dalphin JC, Depner M, Genuneit J, et al. Soluble immunoglobulin A in breast milk is inversely associated with atopic dermatitis at early age: the PASTURE cohort study. Clin Exp Allergy. 2014;44:102–12.

14. Witkowska-Zimny M, Kaminska-El-Hassan E. Cells of human breast milk. Cell Mol Biol Lett. 2017;22:11.

15. Kim JH. Role of Breast-feeding in the Development of Atopic Dermatitis in Early Childhood. Allergy Asthma Immunol Res. 2017;9:285–7.

16. Dawod B, Marshall JS. Cytokines and Soluble Receptors in Breast Milk as Enhancers of Oral Tolerance Development. Front Immunol. 2019;10:16.

17. Järvinen KM, Martin H, Oyoshi MK. Immunomodulatory effects of breast milk on food allergy. Ann Allergy Asthma Immunol. 2019;123:133–43.

18. von Mutius E, Vercelli D. Farm living: effects on childhood asthma and allergy. Nat Rev Immunol. 2010;10:861–8.

19. Ludka-Gaulke T, Ghera P, Waring SC, Keifer M, Seroogy C, Gern JE, et al. Farm exposure in early childhood is associated with a lower risk of severe respiratory illnesses. J Allergy Clin Immunol. 2018;141:454–456.e4.

20. Loss G, Apprich S, Waser M, Kneifel W, Genuneit J, Büchele G, et al. The protective effect of farm milk consumption on childhood asthma and atopy: the GABRIELA study. J Allergy Clin Immunol. 2011;128:766–773.e4.

21. Illi S, Depner M, Genuneit J, Horak E, Loss G, Strunz-Lehner C, et al. Protection from childhood asthma and allergy in Alpine farm environments-the GABRIEL Advanced Studies. J Allergy Clin Immunol. 2012;129:1470–7.e6.

22. Tantoco JC, Elliott Bontrager J, Zhao Q, DeLine J, Seroogy CM. The Amish have decreased asthma and allergic diseases compared with old order Mennonites. Ann Allergy Asthma Immunol. 2018;121:252–253.e1.

23. Holbreich M, Genuneit J, Weber J, Braun-Fahrländer C, Waser M, von Mutius E. Amish children living in northern Indiana have a very low prevalence of allergic sensitization. J Allergy Clin Immunol. 2012;129:1671–3.

24. Stein MM, Hrusch CL, Gozdz J, Igartua C, Pivniouk V, Murray SE, et al. Innate Immunity and Asthma Risk in Amish and Hutterite Farm Children. N Engl J Med. 2016;375:411–21.

25. Martina C, Looney RJ, Marcus C, Allen M, Stahlhut R. Prevalence of allergic disease in Old Order Mennonites in New York. Ann Allergy Asthma Immunol. 2016;117:562–563.e1.

26. Seppo AE, Choudhury R, Pizzarello C, Palli R, Fridy S, Rajani PS, et al. Traditional Farming Lifestyle in Old Older Mennonites Modulates Human Milk Composition. Front Immunol. 2021;12:741513.

27. Steiman CA, Evans MD, Lee KE, Lasarev MR, Gangnon RE, Olson BF, et al. Patterns of farm exposure are associated with reduced incidence of atopic dermatitis in early life [Internet]. Vol. 146, Journal of Allergy and Clinical Immunology. 2020. p. 1379–1386.e6. Available from: 10.1016/j.jaci.2020.06.025

28. Seroogy CM, VanWormer JJ, Olson BF, Evans MD, Johnson T, Cole D, et al. Respiratory health, allergies, and the farm environment: design, methods and enrollment in the observational Wisconsin Infant Study Cohort (WISC): a research proposal. BMC Res Notes. 2019;12:423.

29. Rogier EW, Frantz AL, Bruno MEC, Wedlund L, Cohen DA, Stromberg AJ, et al. Secretory antibodies in breast milk promote long-term intestinal homeostasis by regulating the gut microbiota and host gene expression. Proc Natl Acad Sci U S A. 2014;111:3074–9.

30. Powe CE, Knott CD, Conklin-Brittain N. Infant sex predicts breast milk energy content. Am J Hum Biol. 2010;22:50–4.

31. Fujita M, Roth E, Lo YJ, Hurst C, Vollner J, Kendell A. In poor families, mothers’ milk is richer for daughters than sons: a test of Trivers-Willard hypothesis in agropastoral settlements in Northern Kenya. Am J Phys Anthropol. 2012;149:52–9.

32. Peroni DG, Pescollderungg L, Piacentini GL, Rigotti E, Maselli M, Watschinger K, et al. Immune regulatory cytokines in the milk of lactating women from farming and urban environments. Pediatr Allergy Immunol. 2010;21:977–82.

33. Tomicić S, Johansson G, Voor T, Björkstén B, Böttcher MF, Jenmalm MC. Breast milk cytokine and IgA composition differ in Estonian and Swedish mothers-relationship to microbial pressure and infant allergy. Pediatr Res. 2010;68:330–4.

34. Field CJ. The immunological components of human milk and their effect on immune development in infants. J Nutr. 2005;135:1–4.

35. Toscano M, De Grandi R, Grossi E, Drago L. Role of the Human Breast Milk-Associated Microbiota on the Newborns’ Immune System: A Mini Review. Front Microbiol. 2017;8:2100.

36. Ober C, Sperling AI, von Mutius E, Vercelli D. Immune development and environment: lessons from Amish and Hutterite children. Curr Opin Immunol. 2017;48:51–60.

37. Kirjavainen PV, Karvonen AM, Adams RI, Täubel M, Roponen M, Tuoresmäki P, et al. Farm-like indoor microbiota in non-farm homes protects children from asthma development. Nat Med. 2019;25:1089–95.

38. Medzhitov R. Recognition of microorganisms and activation of the immune response. Nature. 2007;449:819–26.

39. Skovbjerg S, Martner A, Hynsjö L, Hessle C, Olsen I, Dewhirst FE, et al. Gram-positive and gram-negative bacteria induce different patterns of cytokine production in human mononuclear cells irrespective of taxonomic relatedness. J Interferon Cytokine Res. 2010;30:23–32.

40. Tietze K, Dalpke A, Morath S, Mutters R, Heeg K, Nonnenmacher C. Differences in innate immune responses upon stimulation with gram-positive and gram-negative bacteria. J Periodontal Res. 2006;41:447–54.

41. Lahdenperä M, Galante L, Gonzales-Inca C, Vahtera J, Pentti J, Rautava S, et al. Residential green environments are associated with human milk oligosaccharide diversity and composition. Sci Rep. 2023;13:216.

42. Gómez-Gallego C, Morales JM, Monleón D, du Toit E, Kumar H, Linderborg KM, et al. Human Breast Milk NMR Metabolomic Profile across Specific Geographical Locations and Its Association with the Milk Microbiota. Nutrients [Internet]. 2018;10. Available from: 10.3390/nu10101355

43. Kumar H, du Toit E, Kulkarni A, Aakko J, Linderborg KM, Zhang Y, et al. Distinct Patterns in Human Milk Microbiota and Fatty Acid Profiles Across Specific Geographic Locations. Front Microbiol. 2016;7:1619.

44. Fehr K, Moossavi S, Sbihi H, Boutin RCT, Bode L, Robertson B, et al. Breastmilk Feeding Practices Are Associated with the Co-Occurrence of Bacteria in Mothers’ Milk and the Infant Gut: the CHILD Cohort Study. Cell Host Microbe. 2020;28:285–297.e4.

45. Tobin NH, Woodward C, Zabih S, Lee DJ, Li F, Aldrovandi GM. A Method for Targeted 16S Sequencing of Human Milk Samples. J Vis Exp [Internet]. 2018; Available from: 10.3791/56974

46. Assarsson E, Lundberg M, Holmquist G, Björkesten J, Thorsen SB, Ekman D, et al. Homogenous 96-plex PEA immunoassay exhibiting high sensitivity, specificity, and excellent scalability. PLoS One. 2014;9:e95192.

47. Fujimura KE, Sitarik AR, Havstad S, Lin DL, Levan S, Fadrosh D, et al. Neonatal gut microbiota associates with childhood multisensitized atopy and T cell differentiation. Nat Med. 2016;22:1187–91.

48. Caporaso JG, Lauber CL, Walters WA, Berg-Lyons D, Huntley J, Fierer N, et al. Ultra-high-throughput microbial community analysis on the Illumina HiSeq and MiSeq platforms. ISME J. 2012;6:1621–4.

49. Bolyen E, Rideout JR, Dillon MR, Bokulich NA, Abnet CC, Al-Ghalith GA, et al. Reproducible, interactive, scalable and extensible microbiome data science using QIIME 2. Nat Biotechnol. 2019;37:852–7.

50. Callahan BJ, McMurdie PJ, Rosen MJ, Han AW, Johnson AJA, Holmes SP. DADA2: High-resolution sample inference from Illumina amplicon data. Nat Methods. 2016;13:581–3.

51. McDonald D, Jiang Y, Balaban M, Cantrell K, Zhu Q, Gonzalez A, et al. Greengenes2 unifies microbial data in a single reference tree. Nat Biotechnol [Internet]. 2023; Available from: 10.1038/s41587-023-01845-1

52. Austin GI, Park H, Meydan Y, Seeram D, Sezin T, Lou YC, et al. Contamination source modeling with SCRuB improves cancer phenotype prediction from microbiome data. Nat Biotechnol [Internet]. 2023; Available from: 10.1038/s41587-023-01696-w

53. Mallick H, Rahnavard A, McIver LJ, Ma S, Zhang Y, Nguyen LH, et al. Multivariable association discovery in population-scale meta-omics studies. PLoS Comput Biol. 2021;17:e1009442.

54. Rohart F, Gautier B, Singh A, Lê Cao KA. mixOmics: An R package for ’omics feature selection and multiple data integration. PLoS Comput Biol. 2017;13:e1005752.

55. Paulson JN, Stine OC, Bravo HC, Pop M. Differential abundance analysis for microbial marker-gene surveys. Nat Methods. 2013;10:1200–2.

56. Breuer K, Foroushani AK, Laird MR, Chen C, Sribnaia A, Lo R, et al. InnateDB: systems biology of innate immunity and beyond—recent updates and continuing curation. Nucleic Acids Res. 2012;41:D1228–33.

