## Supplementary figures and images for "Farm exposure is associated with human breast milk immune profile and microbiome"

### Supplemental Figure 1

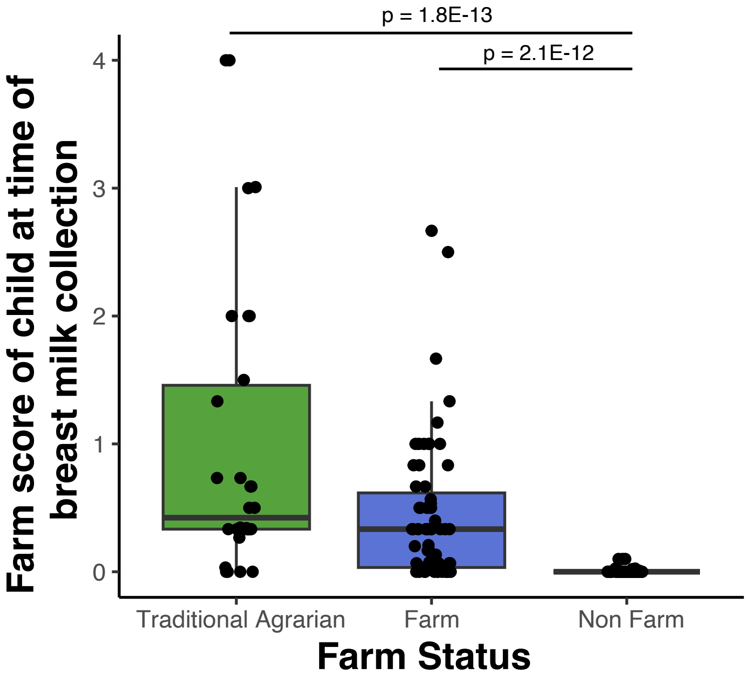

### Supplemental Figure 2

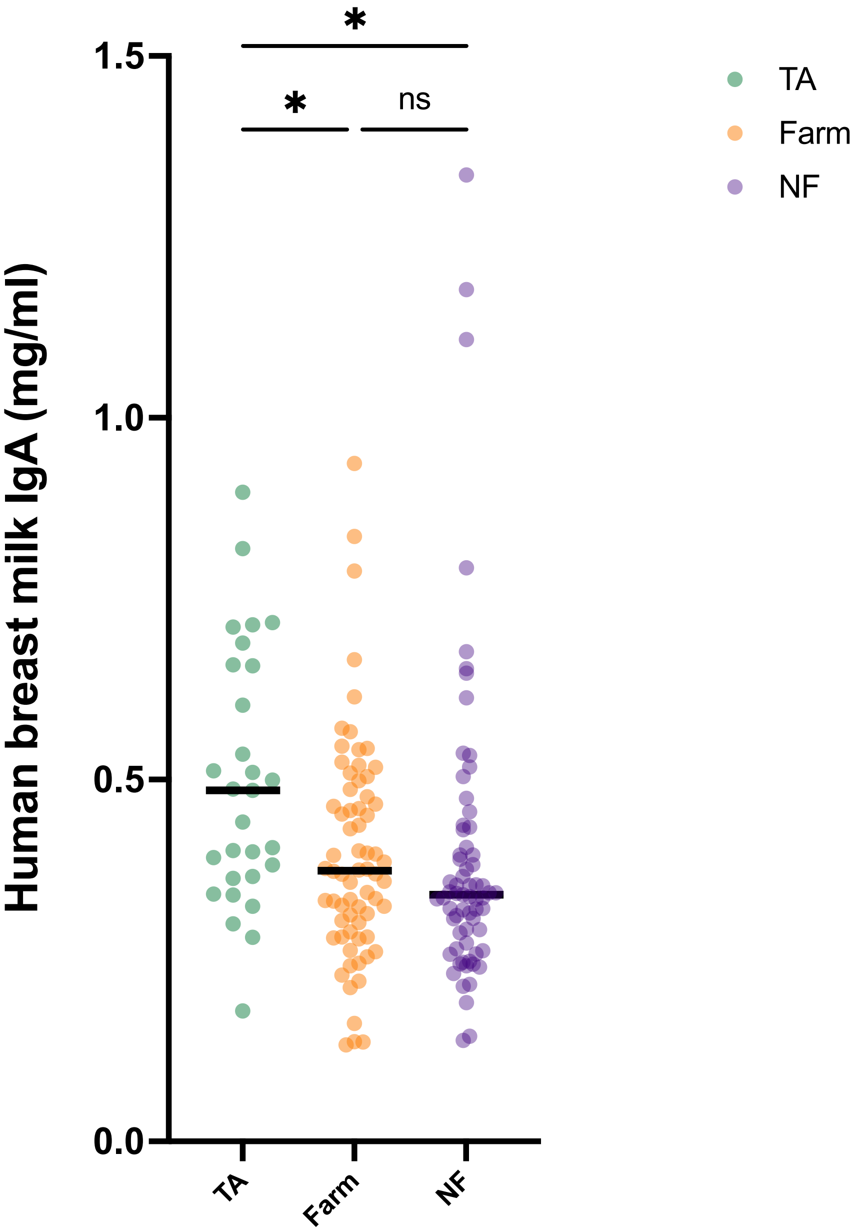

### Supplemental Figure 3

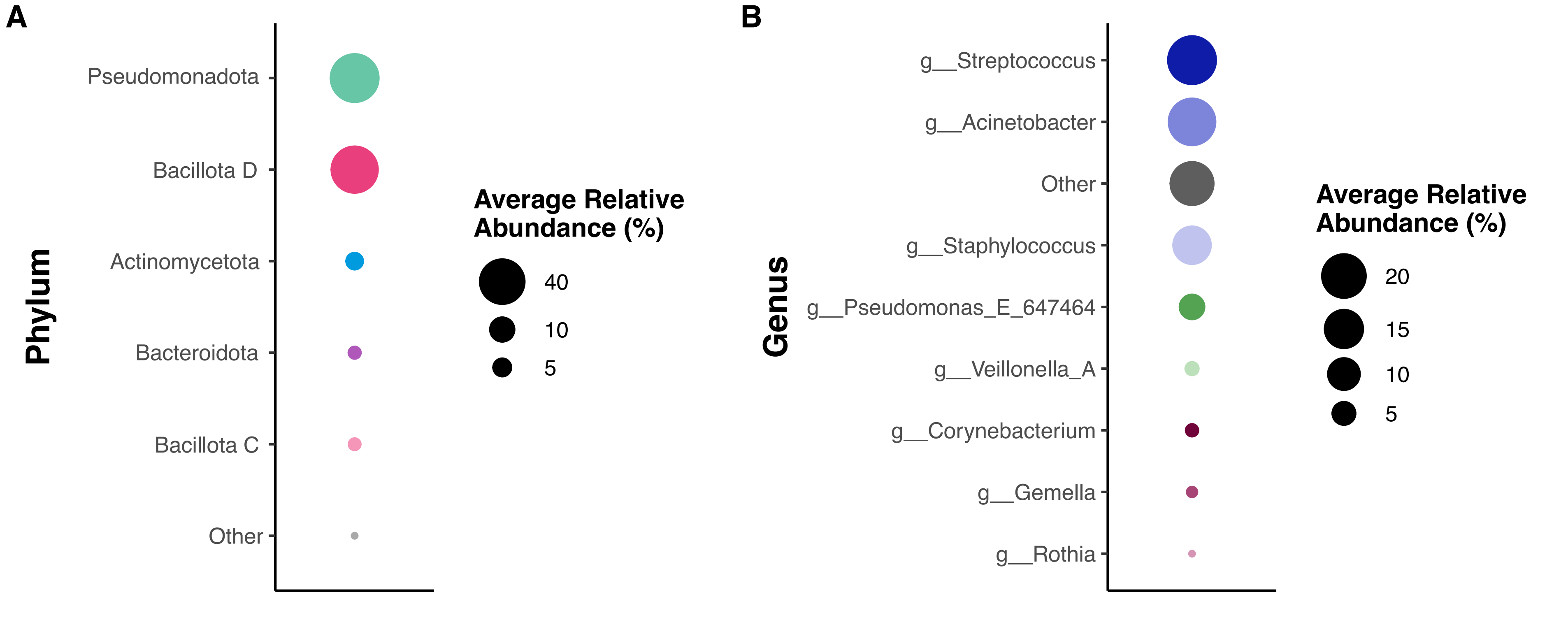

### Supplemental Figure 4

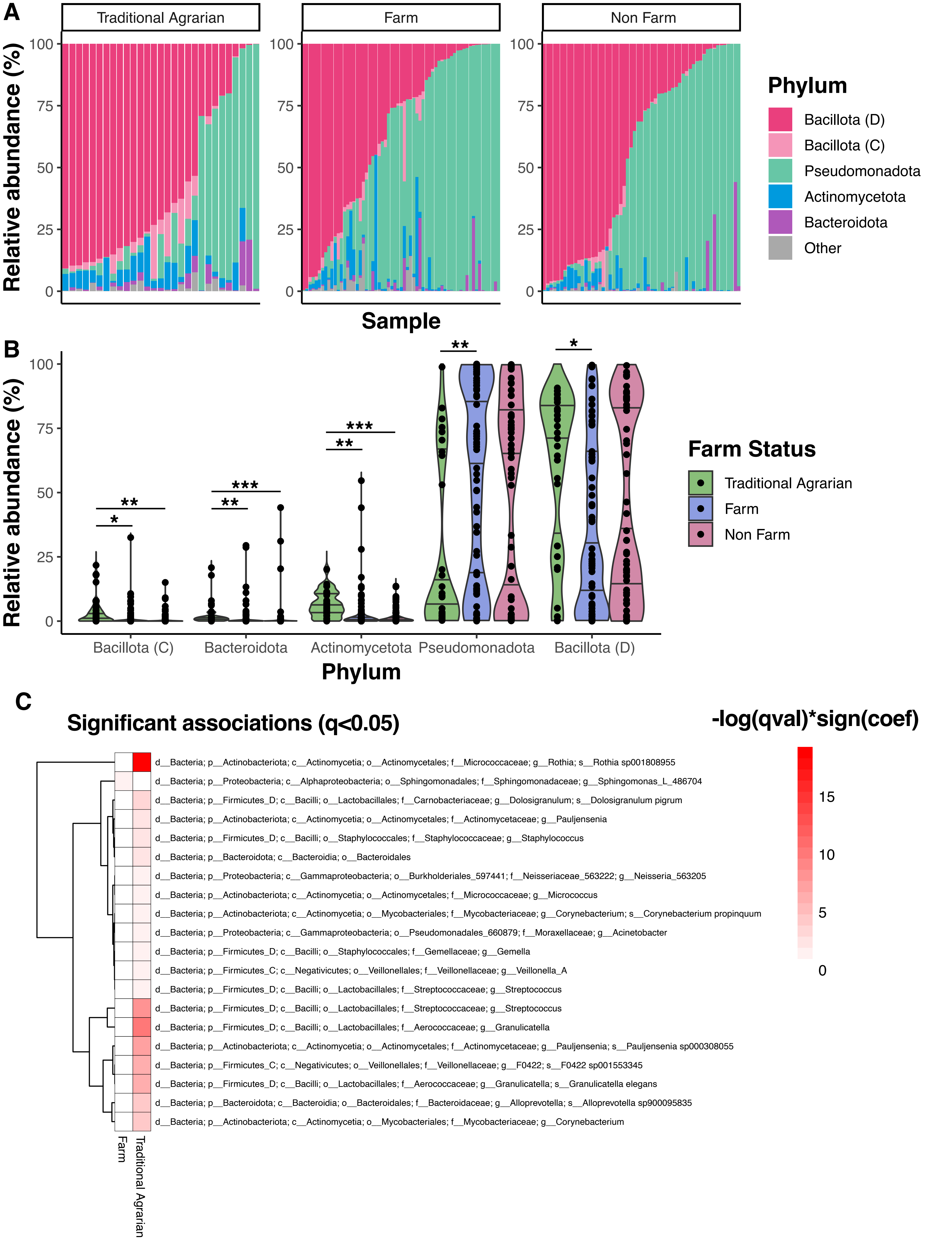
